# Biodetection of an odor signature in white-tailed deer associated with infection by chronic wasting disease prions

**DOI:** 10.1101/2024.04.24.590903

**Authors:** Glen J. Golden, Elizabeth A. Ramirez, Hayley N. Stevens, Jennifer Bourbois, Daniel Grove, Richard A. Bowen, Thomas J. DeLiberto, Bruce A. Kimball

## Abstract

Chronic wasting disease (CWD) has become a major concern among those involved in managing wild and captive cervid populations. CWD is a fatal, highly transmissible spongiform encephalopathy caused by an abnormally folded protein, called a prion. Prions are present in a number of tissues, including feces and urine in CWD infected animals, suggesting multiple modes of transmission, including animal-to-animal, environmental, and by fomite. CWD management is complicated by the lack of practical, non-invasive, live-animal screening tests. We previously demonstrated that ferrets were able to detect and discriminate fecal samples from ducks infected or not infected with avian influenza virus based on odor. That work clearly predicted that animal biodetectors can be trained to identify populations and/or individuals infected with CWD via detection of fecal odors. Such a tool may also prove useful in identifying potentially infected live animals, carcasses, urine, feces, and contaminated environments. Toward this goal, dogs were trained to detect and discriminate CWD infected individuals from non-infected deer in a laboratory setting. Dogs were tested with novel panels of fecal samples demonstrating the dogs’ ability to generalize a learned odor profile to novel odor samples based on infection status. Additionally, dogs were transitioned from alerting to fecal samples to an odor profile that consisted of CWD infection status with a different odor background using different sections of gastrointestinal tracts. These results indicated that canine biodetectors can discriminate the specific odors emitted from the feces of non-infected versus CWD infected white-tailed deer as well as generalizing the learned response to other tissues collected from infected individuals. These findings suggest that the health status of wild and farmed cervids can be evaluated non-invasively for CWD infection via monitoring of volatile metabolites thereby providing an effective tool for rapid CWD surveillance.

## Introduction

It has long been speculated that diseases might be diagnosed based on odor. In the past decade, examination of fecal volatiles has been featured for detection of chronic wasting disease (CWD) in white-tailed deer (*Odocoileus virginianus*) [1], avian influenza virus (AIV) in mallards (*Anas platyrhynchos*) [2, 3]; and bovine tuberculosis in goats (*Capra hircus*) [4, 5], white-tailed deer(WTD) [6], and cattle (*Bos taurus*) [7]. Diagnosis of disease or infection using volatile metabolites found in breath [4, 7–10] or other emanations [11–13] have also been demonstrated. Early research in a mouse model [14] established that MHC genotypes can be discriminated based on whole-body odor odor . Importantly, this same ability to discriminate genotypes was then demonstrated using only the urine collected from live mice (*Mus musculus*) [15]. More recently, studies using both analytical (GC/MS) and biodetector techniques have demonstrated that urine volatile metabolites are altered by immunization [16], injury [17], and inflammation [18]. These and other studies have led researchers to conclude that excreted metabolites can serve as indicators of changes in an animal’s physiological state.

CWD is a fatal, highly transmissible spongiform encephalopathy caused by an abnormally folded protein, called a prion that affects members of the deer family. CWD is a critical management concern in both wild and captive cervid populations. The prion gene, *Prnp*, and the normal prion protein (PrP^C^) are both highly conserved among mammals [19]. Studies using knockout mice and cattle and the naturally occurring absence of the gene in some breeds of goats indicated that the gene is not required for survival and to date the precise function of the prion protein is unknown [19].

Prions are present in a number of tissues, including feces and urine in CWD infected animals, suggesting multiple modes of transmission, including animal-to-animal, environmental, and fomite. CWD is expanding its geographic range through two distinct populations: 1) a more rapid spread through captive herds although this had been slowed by import restrictions if cervids across state lines, and 2) slower through natural movements in free-ranging herds. The clinical syndrome was first recognized in 1967 at the northeastern Colorado/southeastern Wyoming border. CWD has now been detected in free-ranging and captive cervids in 32 states and 5 Canadian provinces. It has also been found outside the North American Continent in Asia and Europe with multiple, separate foci identified [19, 20]

CWD management is complicated by the lack of a practical, non-invasive, live-animal screening test. With the onset of the recent COVID-19 pandemic, there has been an expansion in the use of canine biodetectors in the detection of disease, often without a clear understanding of the fundamentals of mammalian olfaction. Through our work with mice, ferrets (*Mustela furo*), and now dogs (*Canis lupus familiaris*), we have established an understanding of the variables contributing to successful and unsuccessful mammalian biodetection of disease.

A recent study showed that volatile metabolites identified by gas chromatography/mass spectrometry (GC/MS) in feces can be used to discriminate between CWD-positive and -exposed cervids and disease-free cervids [21]. This suggests that volatile metabolites existing in many fluids or tissues could be used to identify CWD infected herds, individuals, carcasses, and environments by trained biodetectors via odor stimuli. The ability for a biodetector to generalize an odor profile from one or a few tissue types and expand it to a variety of tissue/emanations would be extraordinarily valuable. Antemortem methods currently being developed for the detection of CWD are generally not rapid and/or have varying degrees of applicability to different species. Most methods currently used to detect CWD in both farmed and free-ranging cervid populations are conducted postmortem and developing antemortem methods require sedation or restraint for tissue collection [19]. Trained canine biodetectors would provide a tool for collecting information regarding infection status of both farmed and wild populations without the need for either sedation or restraint. Additionally, canine biodetectors would provide a supplemental postmortem testing method that would augment methods already in use. Canine biodetectors that can learn the odor profile of a single tissue type from an infected individual, may also provide a method for detecting subclinical infections earlier than can be achieved by currently accepted methods and may reliably detect CWD prion contaminated environments.

Animal biodetectors have convincingly demonstrated not only that disease diagnoses may be made based on odor alteration [22–24], but that the underlying volatile signals can be detected in the face of environmental variation [25]. Based on the results in mice, we recently trained ferrets to detect avian influenza virus (AIV) infection using fecal volatiles [2]. Not only did trained ferret biodetectors demonstrate that fecal odor was altered in waterfowl by AIV infection, but also that alteration of fecal volatiles was evident to the ferrets before viral shedding occurred with very high sensitivity and specificity. Based on what was learned from training ferrets, dogs (n = 6) were trained in the detection of fecal samples collected from AIV infected ducks. These same dogs can now identify AIV infection in waterfowl via odors of feces, isolated gastrointestinal tracts, and whole duck carcasses (unpublished results).

Dogs were trained to detect and discriminate CWD-infected from non-infected WTD fecal samples in a laboratory setting and were tested in their ability to generalize what was learned from familiar fecal material to novel fecal samples, colon, and novel small intestine samples. This training occurred at the USDA National Wildlife Research Center in Fort Collins, CO using samples from both the Cervid Health Archive (farmed WTD) and samples collected at Ames Plantation in Tennessee (wild WTD). Additionally, field training has been conducted in the CWD-endemic region in southwest Tennessee in which dogs were trained in the busy center of a working farm on Ames Plantation.

## Materials and Methods

### Ethics statement

The experimental protocols for dog research were approved by the National Wildlife Research Center Institutional Animal Care and Use Committee (QA-2870, QA-3155, and QA-3532).

### Biodetectors

Criteria for selecting dogs to serve as potential biodetectors included physical examination, cephalic morphology, and behavioral testing. Once a candidate was identified, other criteria were considered prior to acquisition. This included gender, age, and individual behavioral testing. Dogs were expected to make a discrimination in a complex odor environment that consisted largely of odor cues that will potentially consist of older, decaying sources, as well as fresh fecal samples. Due to these stringent demands, individuals with a strong, enduring motivation to work were considered as having greater potential for success. Six dogs were chosen from shelter and rescue populations from several states and tested for motivation using a modified version of the ball test developed by Working Dogs for Conservation [26].

### White-tailed Deer Feces Stimuli

Fecal samples were obtained from WTD. CWD infection status was identified based on real-time quaking-induced conversion (RT-QuIC), immunohistochemistry (IHC), or enzyme-linked immunosorbent assay (ELISA) from brain tissue, lymph node tissue, or both for confirmation of CWD infection. Samples that were confirmed positive for CWD were categorized as “CWD-positive” and samples in which CWD was not detected were categorized as “CWD-negative”.

### Cohort I

The National Cervid Health Center provided 352 farmed WTD fecal samples collected during targeted removal events conducted throughout the U.S. Information for samples included regional origin, sex, age, codon 96 genotype, and RT-QuIC or IHC results from brain tissue, lymph node tissue, or both for confirmation of CWD infection. Fecal samples (1 g) were placed in1 ml glass vials with screwcap lids (Qorpak, Bridgeville, PA, USA). Sufficient sample material was provided to yield one or more 1 g aliquots.

### Cohort 2

The Tennessee Wildlife Resources Agency (TWRA) provided 166 wild WTD fecal samples collected from hunter harvested animals and animals removed during targeted removal efforts. Samples originated from several counties in western Tennessee. Some of these counties had not yet had a confirmed case of CWD and the others came from areas with reported CWD prevalence ranging from 0.1% to 50%. For example, the number of CWD infected WTD on Ames Plantation brought into the hunter check station was approximately 46%. Sample information included sample origin county and GPS coordinates, sex, age, and CWD infection status determined with CWD ELISA. Fecal samples (1 g) were transferred to 1-mL vials.

### Cohort 3

TWRA provided 81 wild WTD gastrointestinal tract (GI) samples collected from hunter harvested and targeted removal deer. Thirty additional were subsequently provided after experiments had begun. Samples originated from several counties in western Tennessee. Sample information included origin (including GPS coordinates), sex, age, and infection status determined by CWD ELISA. Three or more 1 g aliquots of colon or small intestine were place in 1-mL vials following dissection.

### Presentation of samples and the dogs’ behavioral response

As with the ferrets [27], dogs were trained using 1 in 5 bioassays. Important procedural changes were made to the sample presentation in comparison to the ferrets. Each of the five scratch boxes were attached to the top of one of two stacked 1 ft ^3^ scent boxes (Ray Allen Manufacturing, Colorado Springs, CO, USA) so the scratch box (described below) would be close to nose level for all six dogs. The scent box stacks were spaced 1 ft apart and dogs were trained to use a passive alert (i.e., sit) when detecting CWD-positive fecal samples (Fig 1).

**Fig 1.**
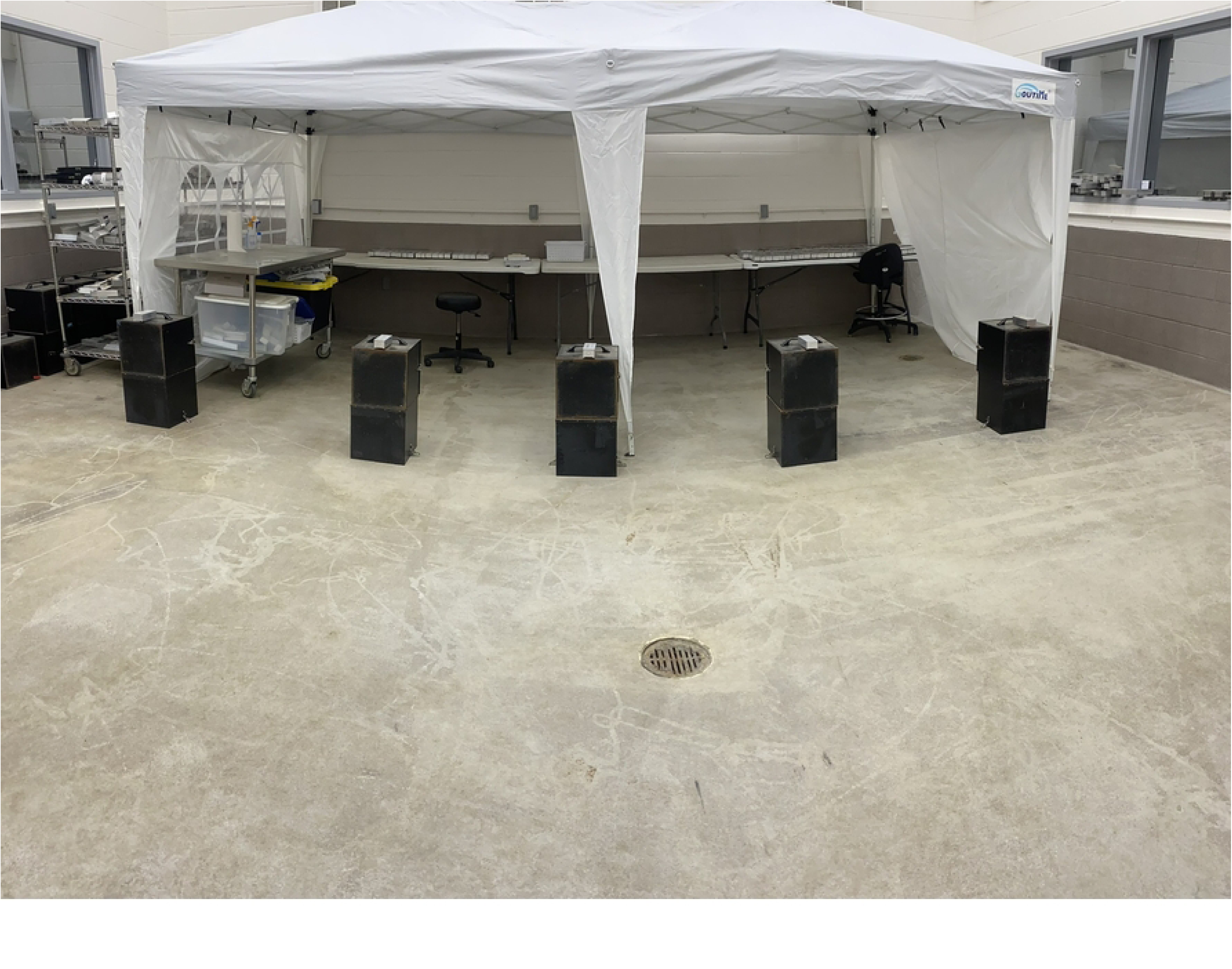
Scratch box assay used to monitor operantly conditioned responses of trained dogs to odors emitted from fecal samples derived from CWD-positive and CWD-negative donor WTD.

**Fig 2.**
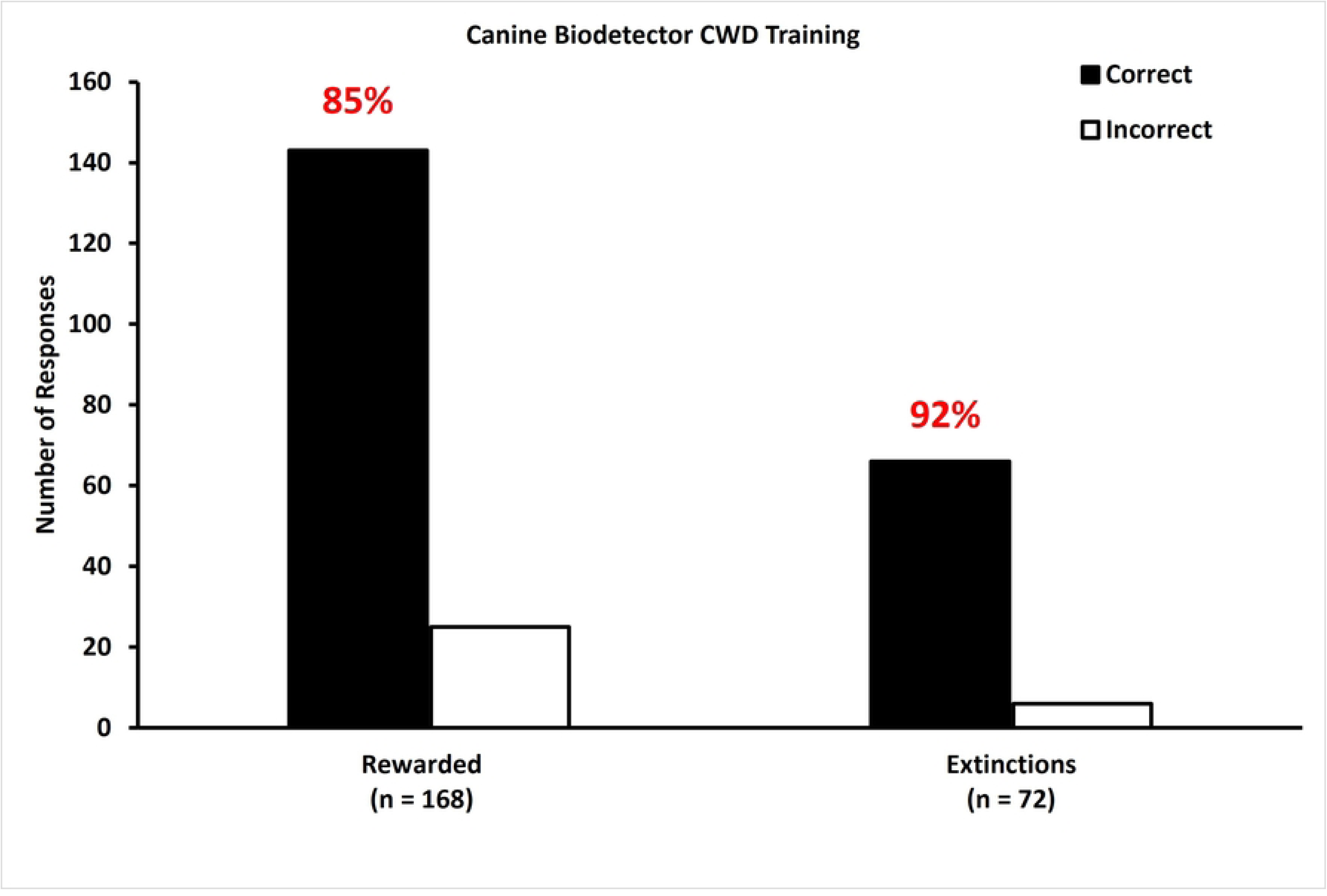
Dogs are accurate in detecting fecal samples from CWD infected WTD during training. During training, dogs correctly (black bars) identified CWD infection in a single fecal sample (n = 240; Cohort 1 and 2) from a CWD infected WTD presented among four CWD-negative samples with greater than 20% accuracy (chance). During rewarded testing trials, dogs correctly identified CWD infection with 85% accuracy. During unrewarded extinction trials, dogs correctly identified infected samples from novel WTD from uninfected samples with 92% accuracy. White bars represent incorrect choices.

A session consisted of 20 trials for each of the six dogs with the position of each box being pseudo-randomized for each trial. During initial training for the passive “sit” response, it took 8 sessions to solidify the desired behavioral response.

For a detailed description of the presentation of fecal samples, please refer to Golden et. al., (2021) [27]. Briefly, the scratch boxes were configured to hold a 1 ml glass vial fitted with filter paper that allowed for the escape of volatiles but not the material placed in the vial (∼1 g of feces per vial). One randomly positioned box of the five held a vial containing fecal material collected from a CWD infected WTD. The remaining four boxes held vials containing fecal material collected from CWD-negative WTD. Feces from an CWD infected WTD were considered the conditioned stimulus positive (CS+) as the dogs were rewarded for alerting to it and feces from CWD-negative WTD were considered the conditioned stimulus negative (CS-).

For the presentation of GI tract samples, the base compartment of each plastic box (16.48 cm x 12.34 cm x 7.6 cm; Glad Products Company, Oakland, CA, USA) was customized with 1” thick foam cut to allow for the retention of a 4 oz glass mason jar (6.00 cm H x 6.35 cm diameter; Homrove, Amazon, USA). Mason jar collar caps (metal septum-type screw caps with a 6.35 cam diameter opening) were fitted with 70 mm, Whatman qualitative filter paper, grade 1 (Sigma-Aldrich, USA) that allowed for the escape of volatiles but not the material placed in the vial (∼1 g of GI colon or small intestine per mason jar). One randomly positioned box of the five held a mason jar containing either a ∼1 g section of colon or small intestine collected from a CWD infected WTD. The remaining four boxes held mason jars containing GI tract tissue collected from CWD-negative WTD. Sections of colon or small intestine from a CWD infected WTD were considered the conditioned stimulus positive (CS+) as the dogs were rewarded for alerting to it and sections of colon or small intestine from CWD-negative WTD were considered the conditioned stimulus negative (CS-).

During training and subsequent testing, a “correct” selection by the dog is the alert given to a 0.01 anethole sample (Table 1; Training 1). When an individual dog correctly alerted to the box containing the CS+ sample, a clicker was activated, and the dog was rewarded with either toys (e.g., tennis or lacrosse balls) or various food treats (e.g., home-made tuna treats, Happy Howie’s) depending on which motivated it best. There was no food restriction and water was available *ad libitum* during training and off-training hours. It is important to note that 20% was the level chance of detecting the CS+ box as only 1 of 5 boxes contained a rewarded sample.

**Table 1.** An overview of the training.

| Experiment | Sample Cohort | Description |
| --- | --- | --- |
| Training 1 | 1 | The passive (sit) alert was shaped in dogs using 0.1M anethole. Dogs were shaped to use a passive alert as a behavioral response to detection of the odor of 0.01M anethole in preparation for work in CWD in 8 sessions. |
| Training 2 | 1 | Dogs were transitioned from the odor of 0.01M anethole to the odor profile of a fecal sample collected from a CWD infected WTD (farmed) by pairing the two odors for one day and then pairing the fecal sample with treats or tennis ball skins in the CS+ box. |
| Training 3 | 1 | Dogs were presented with a panel of five boxes including a fecal sample collected from a CWD infected WTD and four fecal samples collected from CWD-negative WTD. All 20 trials were rewarded with treats or a toy. |
| Training 4 | 1 | Dogs were presented with 20 CS+ and 80 CS- in 20 trials. Once all dogs were performing at 85% or better (chance being 20%), six nonrewarded extinction trials (familiar samples) were introduced to acclimate the dogs to trials without reward. |

### Single-blind procedure

Once dogs proved to be reliable in their response to the CS+, the dogs and the handler were blind to the location of the CS+ to avoid the possibility of the handler inadvertently communicating the position of the CS+ to the dog. An impartial coordinator positioned the CS+ and CS-scratch boxes, signaled the start of a trial, confirmed or rejected the dog handler’s call (described in the next sentence), and picked up the boxes from the previous trial to position the boxes for the next trial. The dog handler controlled when the dogs were to start a trial, called out when a dog alerted to one of the boxes, rewarded the dog if the coordinator confirmed the choice (i.e., clicked the clicker and provided a small food treat or 15 seconds play with a ball or other toy), or without providing verbal or visual cues, turned and walked away if the coordinator called the choice incorrect. The handler and dog were positioned behind a visual barrier prior to the start of each trial as the coordinator prepared the location of the scratch boxes. This procedure was used for all trials that included the shaping of behavior, training, and experimental testing, specifically during rewarded trials but not during unrewarded trials.

### Odor alert response and odor discrimination training

The next three stages of training (Table 1; Training 2-4) please refer to Golden et. al., (2021) While shaping behavior in these early stages, achieving the target of 80% accuracy average for all dogs was considered sufficient and the next training level was initiated.

The following week, the four boxes of PG were replaced with four boxes containing fecal samples collected from CWD-negative WTD. Four fecal samples collected from different CWD-positive WTD donors were used for all 20 rewarded trials. After an initial day of pairing with treats or toys, dogs were trained without pairing for three days. For the paired day, the dogs were able to detect the CS+ with 97% accuracy. For the three unpaired days the dogs were able to detect the CS+ with 88% accuracy (Table 1; Training 3). While shaping behavior in these early stages, achieving the target of 80% accuracy average for all dogs was considered sufficient and the next training level was initiated.

The last stage of training involved using a previously unseen CS+ (n = 20) and four previously unseen CS-(n = 80) fecal samples for each trial. These trials were all rewarded and run in a single-blind procedure as described above. For the first four days, the dogs performed at 73%. The dogs practiced during another 21 days before we introduced six extinction trials. We define an extinction trial as a trial consisting of CS+ and CS-samples that are familiar to the dog (i.e., in terms of the quality, intensity, or donor identity) but is not rewarded or acknowledged in any way. For the last four days of all rewarded, blind trials, the dogs performed at 85%. We then introduced 6 extinction trials out of 20. The dogs performed with 90% accuracy during rewarded trials and 89% accuracy during extinction trials (Table 1; Training 4). While shaping behavior in these early stages, achieving the target of 80% accuracy average for all dogs was considered sufficient and the next training level was initiated.

### Demonstration of learned response

To demonstrate how the trained dogs responded to fecal samples from novel individual CWD infected WTD, we introduced fecal samples from Cohort 2 (wild). For the first two days (Table 2; Demonstration 1, Step 1), the dogs were presented with all rewarded trials. Extinction trials were included for the next two days (Table 2; Demonstration 1, Step 2), with sample panels that were all familiar farmed WTD.

**Table 2.** An overview of the demonstration and experiments.

| Experiment | Sample Cohort | Number of trials | Number of days | Rewarded (Trials; samples) | Extinction (Trials; samples) | Generalization (Trials; samples) |
| --- | --- | --- | --- | --- | --- | --- |
| Demonstration step 1 | 1 and 2 | 20/dog | 2 | 20; familiar mixed | -- | -- |
| Demonstration step 2 | 1 and 2 | 20/dog | 2 | 14; familiar mixed | 6; familiar Cohort 1 | -- |
| Demonstration step 3 | 1 and 2 | 20/dog | 12 | 20; familiar mixed | -- | -- |
| **Preparation for Exp 1** | 1 and 2 | 20/dog | 2 | 14; familiar mixed | 6; familiar Cohort 1 | -- |
| **1** | 1 and 2 | 20/dog | 4 | CS+ 14; mixed<br>CS- 14; Cohort 1 | CS+ 3; familiar mixed<br>CS- 3; familiar Cohort 1 | CS+ 3; novel Cohort 1<br>CS- 3; novel Cohort 1 |
| **2** | 3 | 20/dog | 6 | CS+ 14; mixed familiar colon and small intestine<br>CS- 14; mixed familiar colon and small intestine | CS+ 3; mixed familiar colon and small intestine<br>CS- 3; mixed familiar colon and small intestine | CS+ 3; novel small intestine<br>CS- 3; novel small intestine |

After 12 practice days of all rewarded trials with a mix of both fecal sample Cohorts (Table 2; Demonstration 1, Step 3), extinction trials were included to prepare for experiment 1. For two days, dogs were presented with a mix of both Cohorts in the rewarded trials. For the extinction trials, dogs were presented with panels that consisted of Cohort 2 samples for the four CS- samples and a CS+ from Cohort 1.

### Experiment 1 - Discrimination of feces from CWD infected WTD

The dogs’ responses to samples collected from novel WTD was examined in unrewarded generalization trials interspersed among unrewarded extinction and rewarded trials in four testing sessions on consecutive days (Table 2, Experiment 1). We defined non-rewarded generalization trials here as a trial consisting of CS+ and CS-samples that are novel to the dog (i.e., in terms of the quality, intensity, or donor identity) but is not rewarded or acknowledged in any way. In each session, there were 14 rewarded trials, 3 unrewarded extinction trials and 3 unrewarded generalization trials. There had to be at least 2 rewarded trials prior to either type of unrewarded trial to begin a session, at least 1 rewarded trial between unrewarded trials, and at least one rewarded trial to end the session. Rewarded trial CS+ samples consisted of randomly chosen familiar individual CWD-positive WTD from Cohorts 1 and 2. Rewarded trial CS- samples consisted of randomly chosen familiar individual CWD-negative WTD from Cohort 1.

Extinction trial CS+ samples (3 per session) consisted of randomly chosen familiar individual CWD-positive WTD from Cohort 1. Extinction trial CS- samples consisted of randomly chosen familiar individual CWD-negative WTD from Cohort 2. Generalization trial CS+ samples (3 per session) consisted of randomly chosen novel individual CWD-positive WTD from Cohort 1. Generalization trial CS- samples consisted of randomly chosen novel individual CWD-negative WTD from Cohort 1.

### Experiment 2 – Transition to detection of CWD-positive volatile odors in tissue samples

To determine if dogs can be trained to generalize responses from the odor profile of feces to the odor profile of a GI tissue sample from the same animal, dogs previously trained with defecated feces were also trained with samples of excised colon (approximately 1g and containing feces) in all rewarded generalization trials consisting of a CS+ and four CS- samples and similarly tested with unfamiliar samples not offered during training.

Six days of testing sessions were conducted over two weeks. In each session, there were 14 rewarded trials, 3 unrewarded extinction trials and 3 unrewarded generalization trials. There were at least 2 rewarded trials prior to either type of unrewarded trial to begin a session, at least 1 rewarded trial between unrewarded trials, and at least on rewarded trial to end the session. The reward trials were a mixture of colons and small intestines tissues collected from familiar donors. The extinction trials were familiar colon and small intestine samples. The generalization trials consisted of an entire panel of novel small intestine samples.

### Data Analysis

Cumulative responses across all trained dog trials were calculated for each set of experimental generalization trials. Success rates (number of correct trials divided by the total number of generalization trials) were statistically evaluated using binomial proportion tests with a continuity correction for small numbers of observations [28]. The data were tested for independence between donor identity, testing day, and correct responses using the Cochran-Mantel-Haenszel test [29]. For all experiments, a success rate of 20% would be expected by chance. However, as the goal of these trials was to demonstrate the high specificity of trained biodetectors, dog responses were also compared to 50% and 80% success rates.

## Results

### Demonstration of learned response

To train dogs to respond to fecal samples from novel individual CWD infected WTD and to demonstrate real-world detection, we introduced fecal samples from Cohort 2 (wild). For the first two days of all rewarded trials, the dog’s performance fell to 78% accuracy (from 87% training accuracy with farmed WTD alone). After adding extinctions for two days, which were all familiar farmed WTD, the dogs performed at 90% accuracy during extinctions, but at 80% accuracy during the rewarded trials which were a mix of Cohorts 1 and 2.

After 12 practice days of all rewarded trials with a mix of both Cohorts, extinction trials were added to prepare for experiment 1. For two days, dogs were presented with a mix of both Cohorts in the rewarded trials. For the extinction trials, dogs were presented with panels that consisted of Cohort 2 samples for the four CS- samples and a CS+ from Cohort 1. Dogs performed at 85% accuracy with an average sensitivity of 87% and an average specificity of 96% for the rewarded trials. Dogs performed at 92% accuracy with an average sensitivity of 92% and an average specificity of 98% for the extinction trials.

### Experiment 1 - Discrimination of feces from CWD infected WTD

Trained dogs were highly accurate (85% correct choices in unrewarded generalization trials over 4 testing days) at discriminating between fecal samples collected from CWD-negative WTD and from CWD-positive WTD. (Fig 3).

**Fig 3.**
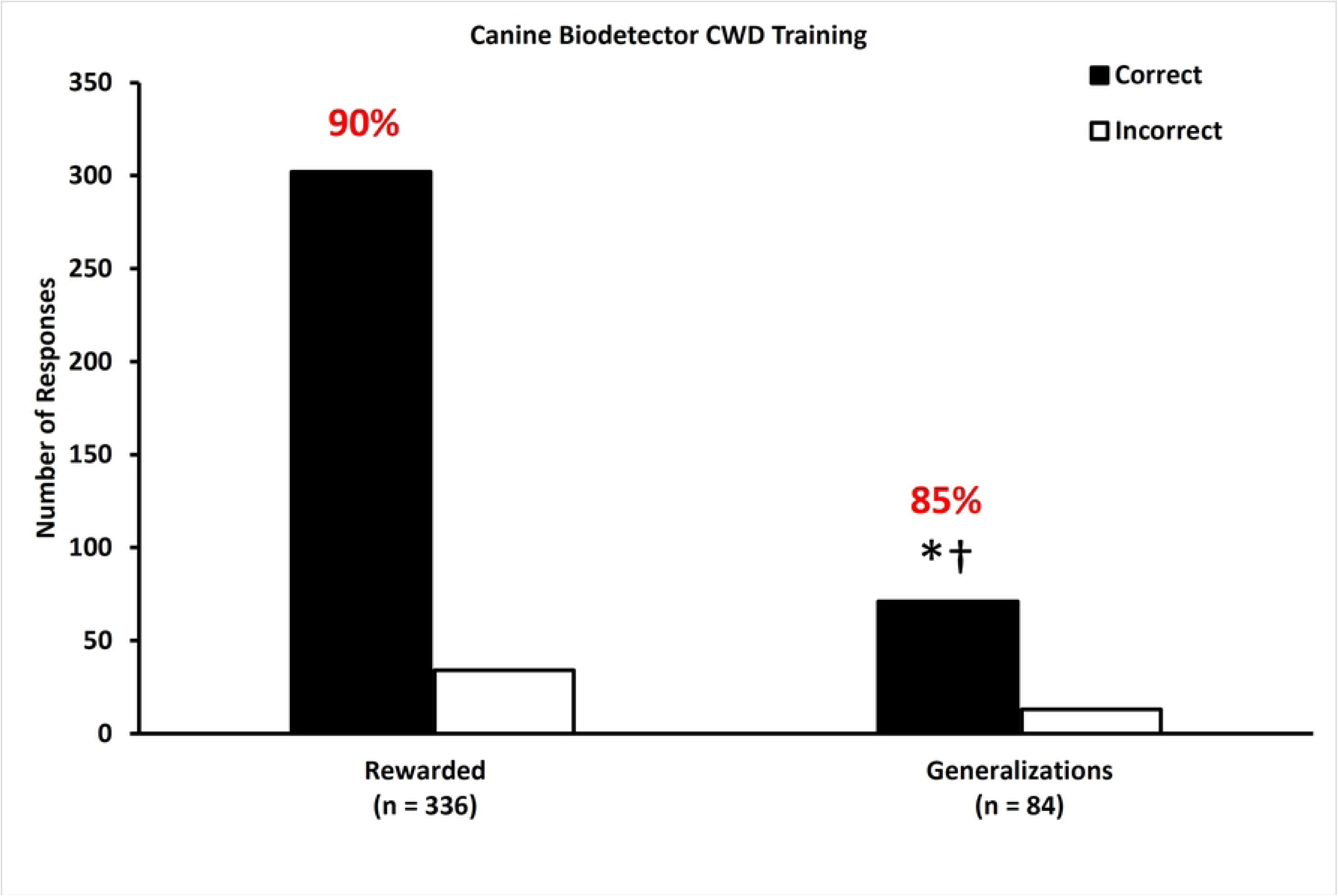
Dogs are capable of generalizing their ability to discriminate between CWD-negative and CWD-positive fecal samples when they encounter fecal samples from novel individuals. During training, dogs correctly (black bars) identified CWD infection in a single fecal sample from a CWD-positive WTD presented among four CWD-negative samples with 85% accuracy. During rewarded testing trials, dogs correctly identified CWD infection with 90% accuracy. During unrewarded generalization trials, dogs correctly identified CWD-positive samples from novel WTD from CWD-negative samples with 85% accuracy, which was greater than null hypotheses of 20% (*p < 0.0001) and 50% (†p < 0.0001), but not different from 80% (p = 0.1500). White bars represent incorrect choices.

All six trained dogs correctly identified CWD infection with 85% accuracy (Fig 3, middle bars) across four days of rewarded testing trials. During unrewarded generalization trials, dogs correctly identified CWD infection with 85% accuracy (Fig 3, right two bars). This result is different from chance (20%; p < 0.0001) and a 50% success rate (p < 0.0001) and is not different from an 80% success rate (p = 0.15). Non-parametric analysis indicated that detection ability did not vary among dogs, and test day was not associated with the selection made by the dogs in either rewarded testing (p = 0.3562) or unrewarded generalization trials (p = 0.4407).

### Experiment 2 - Transition to detection of CWD-positive volatile odors in tissue samples

All six trained dogs correctly identified CWD infection in colon samples with 96% accuracy and in small intestine samples with 76% accuracy (Fig 4a) over six days of rewarded trials. During unrewarded extinction trials (familiar samples) trained dogs correctly identified CWD infection in colon samples with 94% accuracy and in small intestine samples with 88% accuracy (Fig 4b). During unrewarded generalization trials, dogs correctly identified CWD infection in small intestine samples with 79% accuracy (Fig 4c). This result is different from 20% (p < 0.0001), and 50% (p < 0.0001), but not 80% success rates (p = 0.8597). Non-parametric analysis indicated that detection ability did not vary among dogs, and test day was not associated with the selection made by the dogs (p = 0.8452) in these unrewarded generalization trials (p = 0.4031).

**Fig 4.**
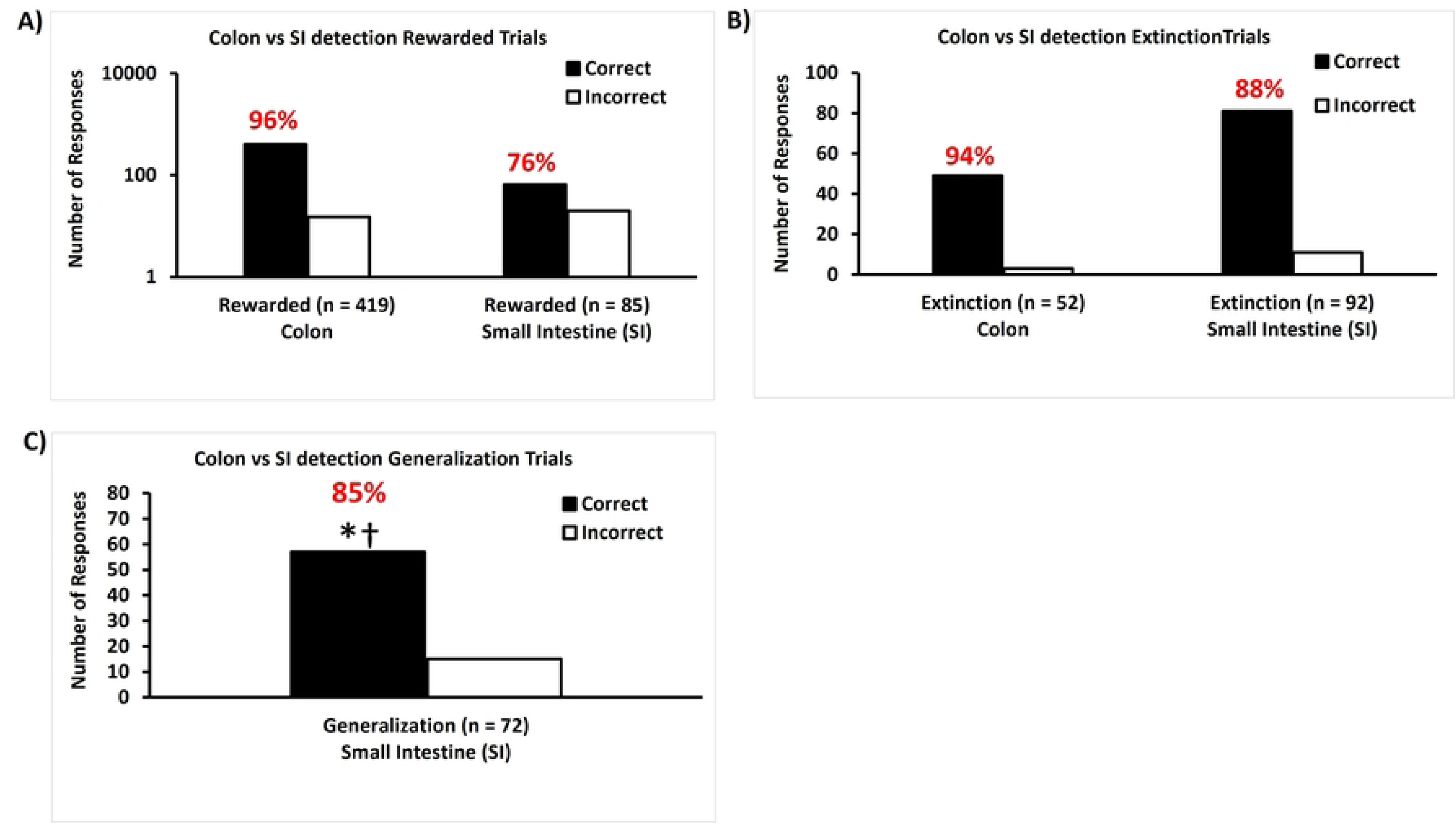
Dogs can identify novel tissue samples from CWD-positive white-tailed deer based on a volatile odor profile that is consistent with a fecal odor profile. A) During 419 rewarded testing trials with a colon sample CS+, dogs correctly (black bars) identified CWD infection with 96% accuracy. During 85 rewarded testing trials with a small intestine sample CS+, dogs correctly (black bars) identified CWD infection with 76% accuracy. B) During 52 extinction testing trials with a colon sample CS+, dogs correctly (black bars) identified CWD infection with 94% accuracy. During 92 extinction testing trials with a small intestine sample CS+, dogs correctly (black bars) identified CWD infection with 88% accuracy. C) During the 72 unrewarded generalization trials, dogs correctly identified CWD infection in small intestine samples with 79% accuracy (*p < 0.0001) in comparison to CWD-negative samples. These classification rates are greater than the null hypotheses of 20% and 50% (†p < 0.0001) success rates, but not for the 80% success rate (p = 0.8597). White bars represent incorrect choices.

## Discussion

This study builds upon previous experiments demonstrating the ability of a mammalian biodetectors to identify disease infection in an animal with high rates of accuracy: e.g. trained mice demonstrated better than 80% accuracy in identifying AIV infection with fecal odors from ducks and ferrets demonstrated better than 90% accuracy for detecting AIV infected ducks [2]. The results of these two experiments provide clear-cut evidence that dogs, are not only capable of performing an olfactory learning discrimination task but can generalize the odor profile learned from fecal samples to other tissues that are not related to the training material.

Importantly, our results suggest that dogs could potentially transition from fecal odor profiles to odor profiles by training them with gastrointestinal tracts dissected into colon and small intestine sections. The colon sections contained fecal material. The small intestine sections did not although they did contain the digested precursor material to feces. Based on the dogs’ response to novel small intestine sections we suggest that dogs can generalize what they learned from the odor profile of fecal samples to other tissues collected from a CWD-positive WTD.

This study echoes previous work demonstrating that dogs can be trained to identify CWD from feces of infected WTD [20]. In that study, Mallikarjun et al. (2023) presented samples on an eight-port scent wheel that became popular with dog cancer detection studies. As in cancer detection studies, the dogs were permitted to explore the wheel without the handler being present allowing for double-blind studies. The current study employed a 1-in-5 bioassay that replicated methods used by the Belgian Police in human scent line-ups [30] such that the same panel could be replicated multiple times with the samples in different positions [31]. In addition, more than three times the number of samples (n = 518; 120 CWD positive and 405 non-infected samples) were presented in the current study. This is important because it has been shown that dogs can easily maintain a working memory of up to 76 odors during an odor memory task [32]. But the most significant aspects of the current study were the testing of GI tract samples following training with fecal samples and the superior CWD sensitivity reported here.

CWD management is complicated by the lack of a practical, rapid, non-invasive, live-animal screening tests. This study adds to our understanding of the fundamentals of mammalian olfaction and how it can be leveraged to be used as an important tool in the surveillance and management of CWD. We have established an understanding of what variables contribute to successful and unsuccessful mammalian biodetection of CWD. Another critical tool would be to determine if dogs can detect the urine and feces of CWD infected deer at historical bait stations and long vacated salt licks. Or the detection of prions at scrapes in high prevalence locations. Another interesting question is do inert prions, free from a host, interact with the natural microbiome of the soil itself producing an odor specific change in the volatile profile of the soil making it possible to detect large deposits of prions?

In the previous ferret study, ferrets could detect the volatile odor profile associated with avian influenza virus infection before the virus was detectable by qPCR [2]. We have no evidence to present that the dog trained to detect the results of CWD infection can do something similar, but it would be very interesting to investigate this question. The farmed samples donated by the National Cervid Health Archive were tested using either lymph node or brain tissue or both. The dogs did not show a difference in their capability to detect any of those samples, but it would beneficial to have fecal samples collected from a staged infection study so the dogs could be challenged with samples from the earliest stages of CWD infection.

The current study provides evidence that biodetectors can be used to detect fecal samples from WTD infected with CWD. This evidence shows that dogs can identify the signature odor that is a result of infection from CWD whether it be in a fecal sample or in a small intestine sample. Regardless, these experiments raised some new questions that need to be addressed. For example, analysis of the data shows that the accuracy of the dogs was diminished by 10% when the wild WTD samples were introduced. This suggests that the volatile odor profile of CWD infected wild WTD may be sufficiently different from farmed CWD infected WTD; or at least different enough to challenge the perception of the volatile odor profile the dogs learned in training on the samples collected from farmed WTD. Most importantly, it will be interesting to determine whether a biodetector trained to identify CWD in WTD fecal samples could identify the odor identity of CWD in another species. This study clearly demonstrates the feasibility of deploying trained biodetectors for CWD surveillance in WTD.

## Acknowledgements

The authors thank Jeremy Dennison and his TWRA staff for assistance in sample collection from wild WTD. Special thanks to Stephanie Karns and Sonia Mongold for being the TWRA liaison and facilitating our unique needs. Special thanks to Amelia Johnston and Jacki Young for some data analysis and canine husbandry. Mention of specific products does not constitute endorsement by Colorado State University, the United States Department of Agriculture or Monell Chemical Senses Center.

